# High-Intensity Interval Training Remodels Adipose Tissue Inflammatory Signaling and Enhances Immunometabolic Health via microRNA Regulation

**DOI:** 10.64898/2026.07.01.735944

**Authors:** Mohammadreza Sadeghi Mohammadi, Sayyed Mohammad Marandi, Zeinab Rezaee, Nicholas J. Saner, Mina Poosti

**Affiliations:** Faculty of Sport Sciences, University of Isfahan, Isfahan, Iran; Institute for Health and Sport, Victoria University, Melbourne, Australia

**Author notes:** **Corresponding Authors:** ➢ Mina Poosti; HDR of Exercise Physiology, University of Isfahan, Isfahan, Iran. ^1^ Present address: Institute for Health and Sport, Victoria University, Melbourne, Australia (Visiting Research Student);, ➢ Sayyed Mohammad Marandi; Professor of Exercise Physiology, University of Isfahan, Isfahan, Iran.

**Keywords:** NLRP3, TNF-α, PPAR-γ, IL-10, miR-21, miR-30d-5p, Wistar rat, Exercise

## Abstract

Sedentary behavior promotes chronic low-grade inflammation in adipose tissue, contributing to metabolic dysfunction and insulin resistance. High-intensity interval training (HIIT) is a time-efficient exercise strategy with potent anti-inflammatory and metabolic benefits; however, its effects on adipose tissue inflammatory signaling and microRNA (miRNA) regulation remain incompletely understood. This study investigated the effects of eight weeks of HIIT on inflammatory and epigenetic markers in interscapular white adipose tissue (iWAT) of male Wistar rats. Fourteen rats were randomly assigned to either a sedentary (SED; n = 7) or HIIT (n = 7) group. The HIIT protocol consisted of treadmill running five days per week for eight weeks. Body weight and iWAT mass were assessed, and molecular adaptations were evaluated at multiple regulatory levels using RT-qPCR for mRNA targets (NLRP3, TNF-α, PPAR-γ, and IL-10) and miRNAs (miR-21 and miR-30d-5p), while protein levels of NLRP3 and PPAR-γ were assessed using Western blotting. Compared with the SED group, HIIT significantly reduced body weight (p < 0.001) and iWAT mass (p = 0.002). Furthermore, HIIT downregulated the expression of pro-inflammatory mediators, including NLRP3 (gene: p = 0.001; protein: p < 0.001) and TNF-α (p = 0.025), while upregulating anti-inflammatory regulators PPAR-γ (gene: p = 0.026; protein: p = 0.020) and IL-10 (p = 0.010). In parallel, inflammation-associated miRNAs, including miR-21 (p = 0.004) and miR-30d-5p (p = 0.002), were markedly downregulated. These coordinated transcriptional, post-transcriptional, and translational adaptations suggest that HIIT attenuates adipose tissue inflammation and promotes a favorable immunometabolic phenotype through integrated molecular and epigenetic mechanisms.

## 1. Introduction

In recent years, the prevalence of sedentary lifestyles have markedly increased and are now recognized as a major contributor to the rising prevalence of metabolic disorders such as obesity, type 2 diabetes, and metabolic syndrome (1, 2). This has been closely linked to chronic low-grade inflammation in key metabolic organs, including liver, skeletal muscle, and adipose tissue (2-4). Among these organs, white adipose tissue (WAT) has emerged as a central regulator of immune and metabolic homeostasis, secreting cytokines and adipokines that can either promote or suppress inflammation. Dysregulation of these mediators under conditions of inactivity or obesity fosters a persistent pro-inflammatory state, impairing insulin signaling and metabolic health (4, 5).

Key molecular regulators of WAT inflammation include peroxisome proliferator-activated receptor gamma (PPAR-γ), which promotes adipocyte function and suppresses inflammatory signaling (6), and NOD-like receptor protein 3 (NLRP3), an inflammasome component that amplifies pro-inflammatory cytokine production, such as interleukin (IL)-1β and IL-18 (7). Moreover, the balance between tumor necrosis factor-alpha (TNF-α), a canonical pro-inflammatory cytokine, and IL-10, a key anti-inflammatory mediator, reflects the immune equilibrium within WAT (8, 9).

Epigenetic mechanisms, particularly microRNAs (miRNAs), have emerged as pivotal regulators of these inflammatory pathways (10). Dysregulation of miRNA expression in WAT has been implicated in obesity, insulin resistance, and type 2 diabetes, highlighting their importance in the development of metabolic disease (11). For instance, miR-21 is often elevated under inflammatory conditions, where it promotes macrophage activation and insulin resistance (11, 12), while miR-30d has been associated with adipocyte differentiation, glucose metabolism, and regulation of inflammatory signaling (13-15). These miRNAs act as molecular intermediates linking environmental stimuli to adaptive metabolic and immune responses in adipose tissue (11-13).

Regular exercise is a potent non-pharmacological intervention that improves metabolic health by reducing adiposity, modulating inflammatory pathways, and enhancing insulin sensitivity (16-18). In particular, moderate-intensity continuous training (MICT) has been widely shown to attenuate adipose tissue inflammation by reducing pro-inflammatory cytokine expression and improving adipokine profiles, thereby contributing to enhanced insulin sensitivity and metabolic homeostasis (19, 20). High-intensity interval training (HIIT), characterized by repeated bouts of vigorous exercise interspersed with active or passive recovery, has emerged as a time-efficient alternative to MICT (21, 22). While current evidence supports that both MICT and HIIT effectively improve body composition, cardiorespiratory fitness (CRF), and promote overall metabolic health (23), recent evidence suggests that HIIT can induces greater improvements in CRF, glucose regulation, and insulin sensitivity compared with MICT (21, 22). Furthermore, the unique physiological stress induced by HIIT may modulate inflammatory signaling pathways differently from traditional endurance exercise (24-27), highlighting the importance of understanding its impacts on inflammatory regulation at the molecular level.

Despite growing interest in HIIT as a potent exercise modality, its effects on the coordinated regulation of inflammatory genes, proteins, and microRNAs (miRNAs) in WAT remains unclear. Therefore, the present study investigated the effects of eight weeks of HIIT on inflammatory signaling in interscapular WAT (iWAT) of male Wistar rats, focusing on the expression of key inflammatory genes and proteins alongside the regulatory miRNAs miR-21 and miR-30d-5p.

## 2. Materials and Methods

### 2.1. Animal Experiment

Fourteen male Wistar rats (8 weeks old), were obtained from the Animal House of Isfahan University of Medical Sciences (Iran, Isfahan). Animals were housed under controlled conditions (temperature 23-24°C, relative humidity 50-53%, and a 12:12 h light-dark cycle) with free access to standard laboratory chow and water. After a 1-week acclimatization period, rats were randomly allocated into two groups (n=7 per group): (1) Sedentary (SED) and (2) high-intensity interval training (HIIT).

All procedures were conducted in accordance with ARRIVE guidelines and the instructions of the ethics committee in research and care of animals of Institute of Sports Sciences of Iran. The study protocol received formal approval from the Ethics Committee of the University of Isfahan IR.UI.REC.1404.192.

### 2.2. Exercise Training Protocol

The HIIT protocol was carried out on a motorized treadmill, five sessions per week for eight weeks, preceded by a one-week familiarization period. Each training session began with a 3-min warm-up and consisted of three repeated exercise bouts. Each bout included a 3-min running period at 15 m/min followed by a 7-min high-intensity interval. The speed of the high intensity intervals was progressively increased from 20 m/min during the first week to 30 m/min by week 8, while treadmill inclination remained at 0% throughout the intervention. Bouts were separated by 2-min recovery periods, and each session ended with a 3-min cool-down (Fig. 1) (28). To ensure adherence, mild electrical stimuli (< 1 mA) were applied only if animals refused to run, and their use was strictly limited to reduce the risk of confounding. The SED group was placed on treadmill without exercise to control for handling-related stress.

**Fig. 1.**
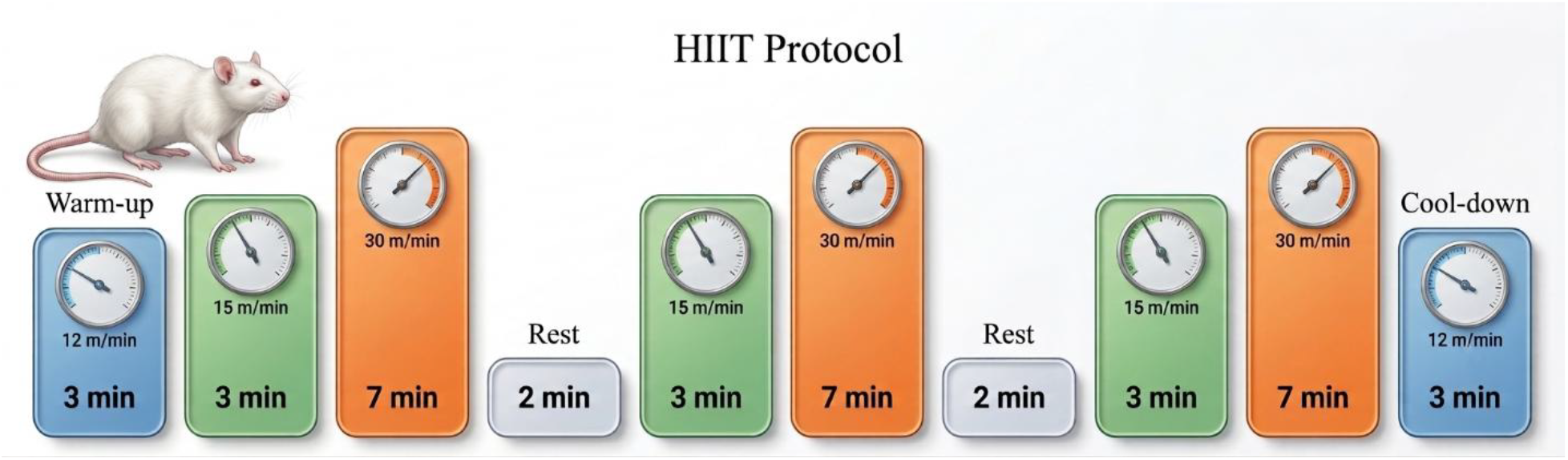
Schematic overview of the High-intensity interval training (HIIIT) protocol and treadmill setup used for rat training. Exercise sessions were performed five days per week, for eight weeks.

### 2.3. Body Weight and Adipose Tissue Mass Assessment

Body weight was measured every two weeks throughout the experimental period. At the time of sacrifice, the weight of iWAT was also measured immediately after excision.

### 2.4. Tissue Collection

Seventy-two hours after the final training session and following an overnight fast, rats were anesthetized by intraperitoneal injection of xylazine (10 mg/kg) and ketamine (80 mg/kg). iWAT was excised from the interscapular region, rinsed in saline, snap-frozen in liquid nitrogen, and stored at -80°C until further molecular analyses.

### 2.5. RNA and microRNA Expression Analyses

Total RNA was isolated from approximately 50 mg of powdered iWAT. For mRNA analysis, RNA extraction was performed using a Total RNA Extraction Kit (Pars Tous, Iran), whereas total RNA including small RNAs was isolated for microRNA analysis using the FavorPrep™ miRNA Isolation Kit (Favogen, South Korea). RNA concentration and purity were assessed spectrophotometrically.

First-strand cDNA for mRNA quantification was synthesized from 1 μg of total RNA using the Easy™ cDNA Synthesis Kit (Pars Tous, Iran) with oligo(dT) and random hexamer primers. For microRNA analysis, cDNA synthesis was performed using stem-loop primers designed based on sequences obtained from the sRNAPrimerDB database.

Quantitative real-time PCR (qRT-PCR) was carried out using SYBR Green Master Mix (Pars Tous, Iran) on a Lava96 Real-Time PCR Detection System (DaAnGene, China). The expression levels of TNF-α, NLRP3, PPAR-γ, and IL-10 were normalized to GAPDH, while miR-21 and miR-30d-5p expression levels were normalized to U6 small nuclear RNA. All reactions were performed in triplicate. Relative expression levels were calculated using the 2^-ΔΔCt^ method. Primer sequences and amplicon characteristics for mRNA and microRNA analyses are provided in Tables 1 and 2, respectively.

**Table 1.**
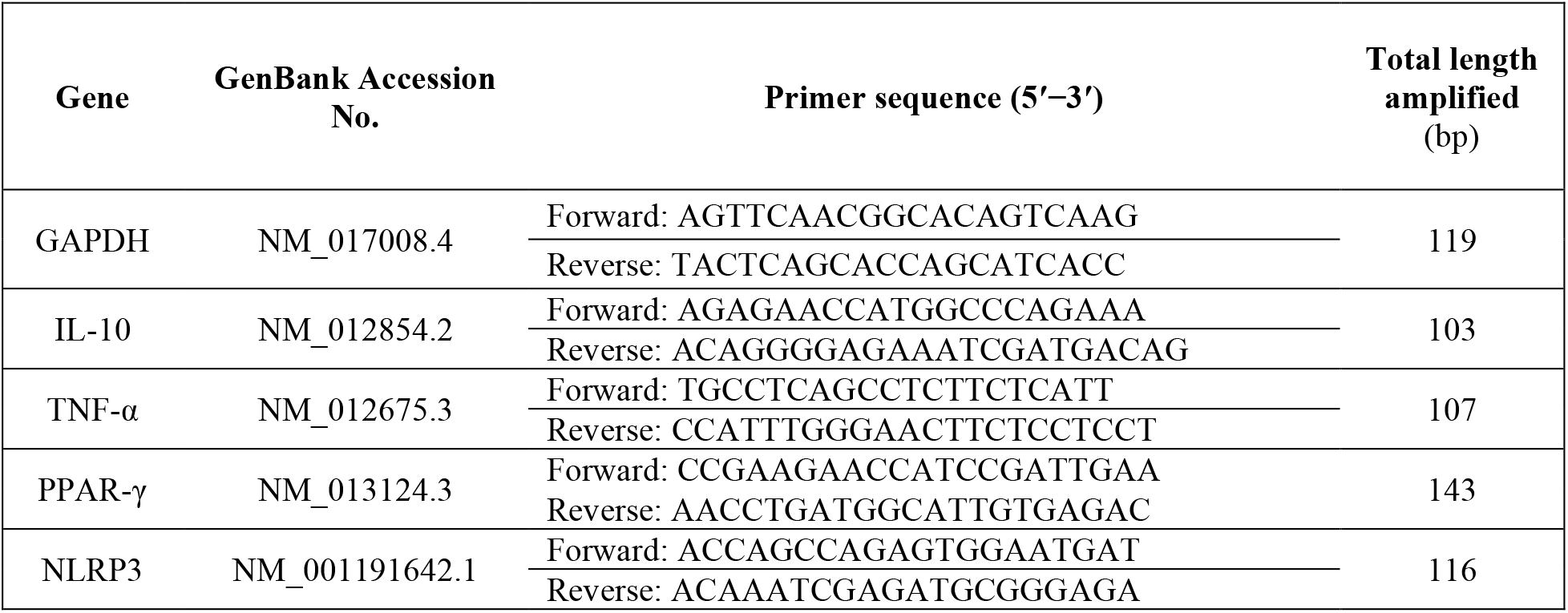
Primers used for mRNA expression analysis in iWAT.

**Table 2.**
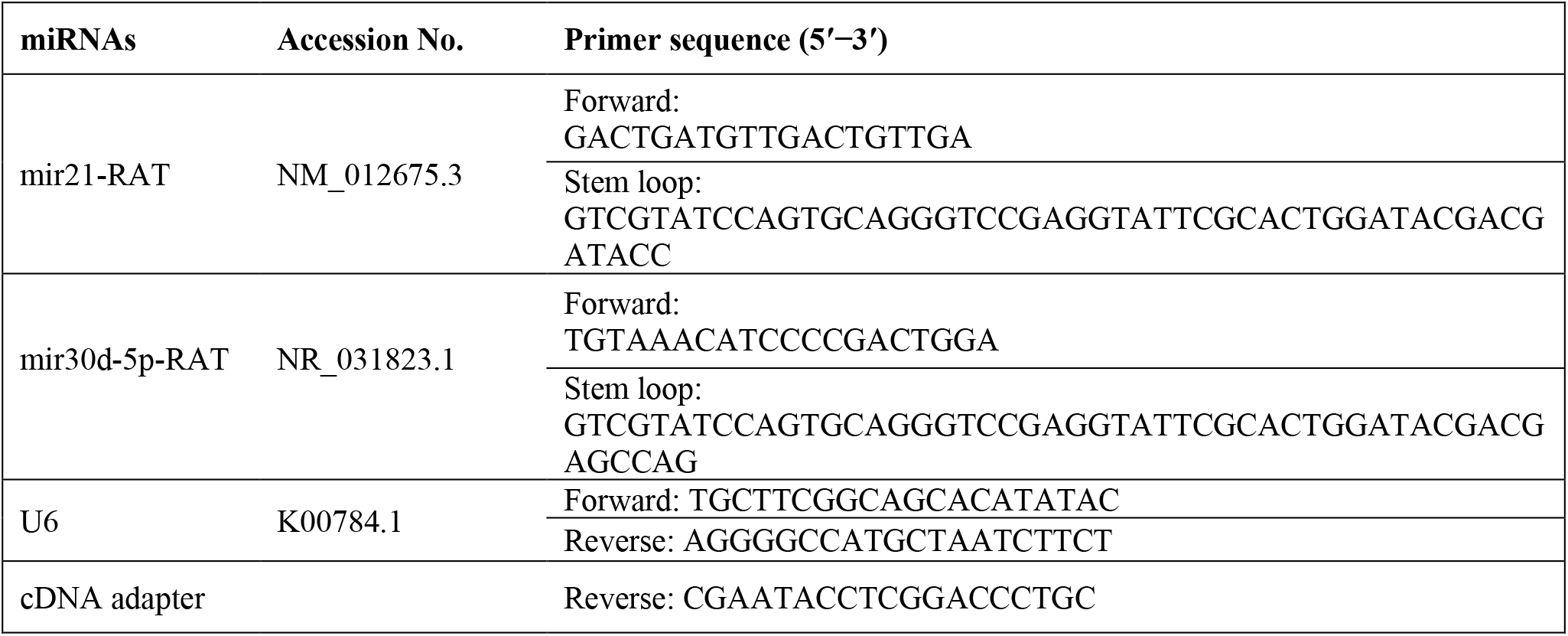
Primers used for microRNA expression analysis in iWAT.

### 2.6. Protein Expression Analysis (Western Blot)

Protein extracts from iWAT were prepared using ice-cold RIPA lysis buffer containing a protease inhibitor cocktail (Sigma, USA). Homogenates were centrifuged at 14,000 rpm for 10 min at 4 °C, and protein concentrations were determined using the Bradford assay with bovine serum albumin as the standard.

Equal amounts of protein were separated by 12% SDS-PAGE and transferred onto PVDF membranes (0.45 μm, Amersham, USA). Membranes were blocked with 5% skim milk and incubated with primary antibodies against PPAR-γ (Cell Signaling Technology, #95128) and NLRP3 (antibodies-online, ABIN1386361). GAPDH (Cell Signaling Technology, #5174) served as the loading control. Protein bands were detected using an enhanced chemiluminescence (ECL) kit (Pars Tous, Iran) and visualized with the FUSION FX imaging system (Vilber, USA). Densitometric analysis was performed using FUSION FX software, and protein expression was normalized to GAPDH.

### 2.7. Statistical Analysis

Data were analyzed using SPSS version 26.0 (IBM Crop., Armonk, NY, USA). Normality was assessed with the Shapiro-Wilk test. Between-group comparisons were performed using independent-sample T-test. Effect size was reported using eta-squared (η^2^) and categorized as small (0.01 ≤ η^2^ < 0.06), medium (0.06 ≤ η^2^ < 0.14), and large (η^2^ ≥ 0.14) . In addition, 95% confidence intervals (95% CI) for the mean differences between groups were calculated and reported (29). All gene, miR and protein expression results were normalized to the SED group. Figures were generated with GraphPad Prism version 10.4.1 (GraphPad Software, San Diego, CA, USA). Results are expressed as mean ± standard deviation (SD), and statistical significance was set at p < 0.05.

## 3. Results

### 3.1. Body Composition

At baseline, body weight did not differ significantly between groups (SED: 246.4 ± 8.2 g vs. HIIT: 247.7 ± 10.9 g, [95% CI -12.55 to 10.01], p = 0.81, η^2^ = 0.005). At the end of the intervention, body weight was significantly lower in the HIIT group compared with the sedentary group (SED: 313.6 ± 29.7 g vs. HIIT:

201.4 ± 36.6, [95% CI -151.1 to -73.43], p < 0.001, η^2^ = 0.768), indicating a regulatory effect of HIIT on body weight (Fig. 2-A).

**Figure 2.**
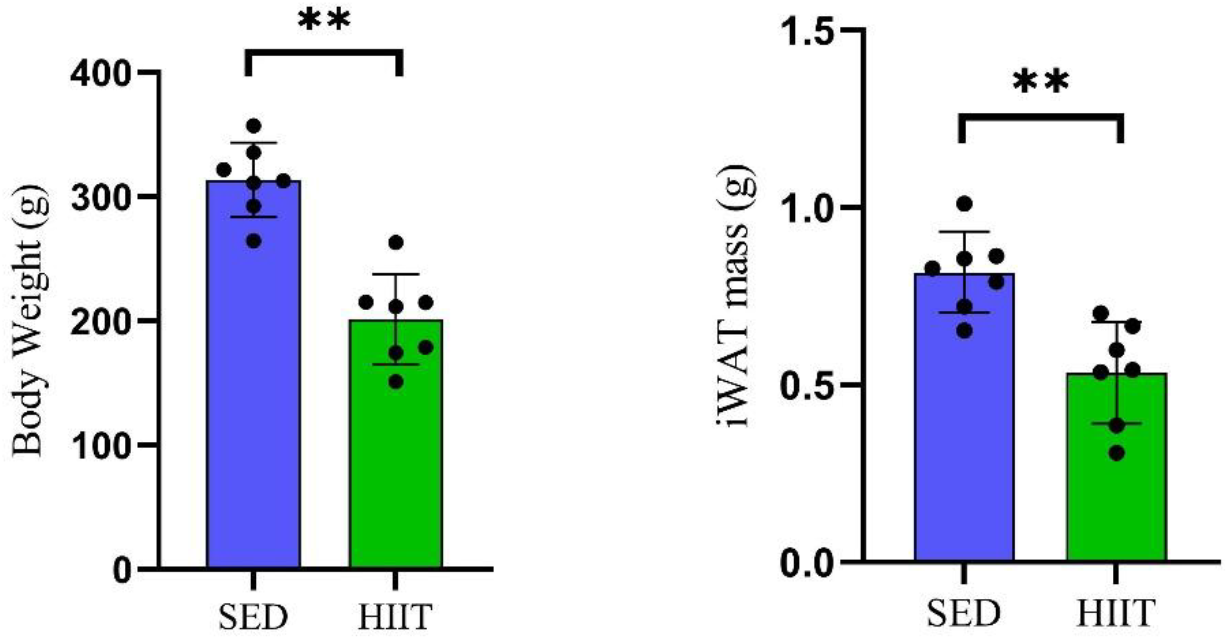
Effects of eight weeks of HIIT on body composition. (A) Body weight. (B) iWAT mass. Data are presented as mean ± SD. SED: sedentary; HIIT: high-intensity interval training. ** p ≤ 0.001 versus SED.

Consistently, iWAT mass was markedly reduced in HIIT group relative to sedentary group (SED: 0.82 ± 0.11 g vs. HIIT: 0.53 ± 0.14 g, [95% CI -0.43 to -0.13], p = 0.002, η^2^ = 0.584), demonstrating effective attenuation of fat accumulation (Fig. 2-B).

### 3.2. Molecular analyses

HIIT modulated inflammatory mediators at multiple molecular levels in a coordinated manner. At the transcriptional level, HIIT significantly reduced NLRP3 gene expression by 58% ([95% CI -0.84 to -0.33], p = 0.001, η^2^ = 0.778) and TNF-α gene expression by 52% ([95% CI -1.02 to -0.09], p = 0.025, η^2^ = 0.486), indicating suppression of pro-inflammatory signaling (Fig. 3-A), In contrast, HIIT increased PPAR-γ gene expression by 171% ([95% CI 0.27 to 3.15], p = 0.026, η^2^ = 0.483) and IL-10 gene expression by 140% ([95% CI 0.46 to 2.50], p = 0.01, η^2^ = 0.585) reflecting enhanced anti-inflammatory responses, compared to the SED group (Fig. 3-B).

**Figure 3.**
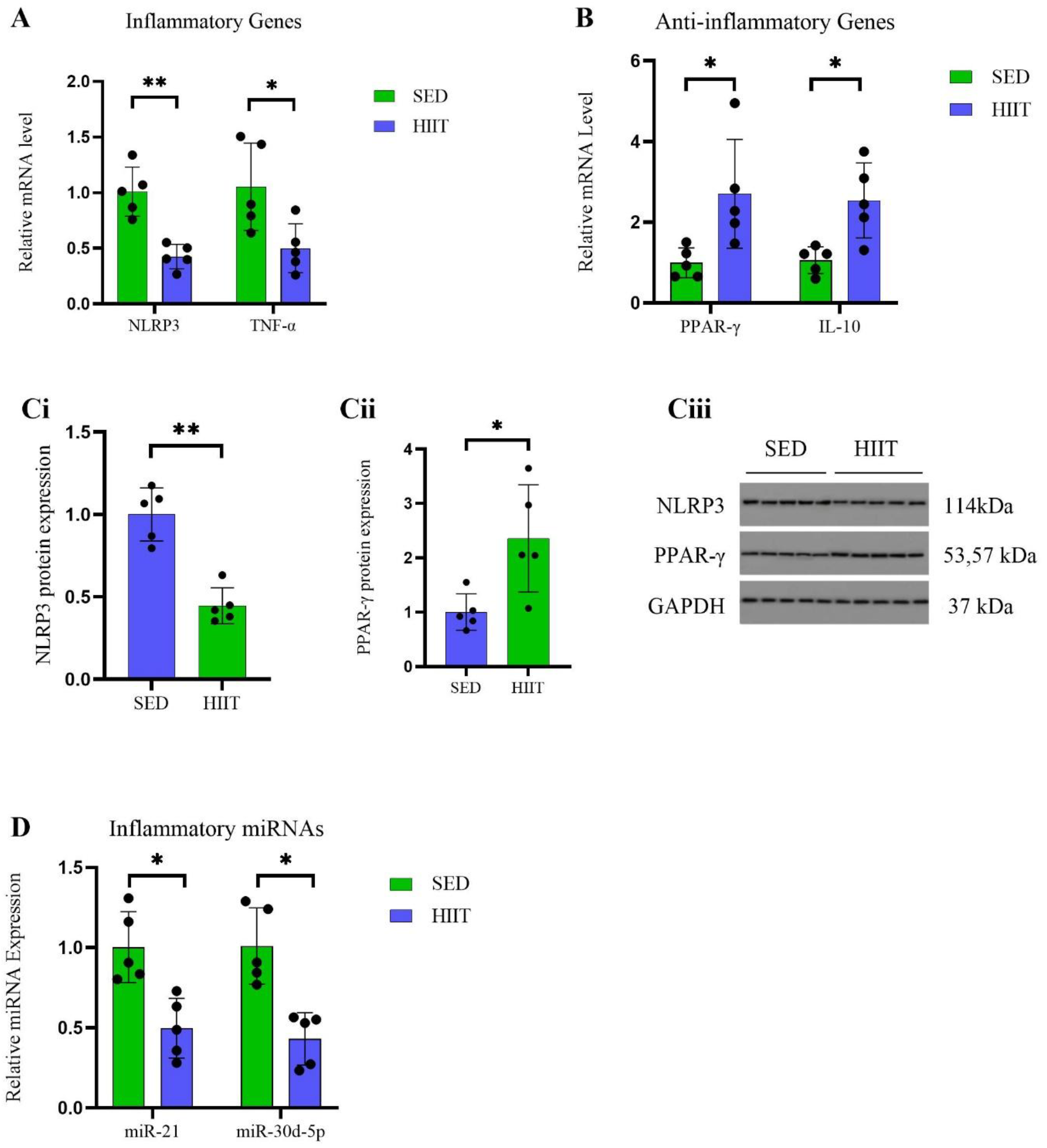
Effects of eight weeks of HIIT on inflammatory signaling, compared to the SED group, in iWAT of male Wistar rats. (A) Relative mRNA expression of pro-inflammatory genes NLRP3 and TNF-α. (B) Relative mRNA expression of anti-inflammatory genes PPAR-γ and IL-10. (Ci) Quantitative densitometry of NLRP3 protein. (Cii) Quantitative densitometry of PPAR-γ protein. (Ciii) Representative Western blot images showing band intensities for NLRP3, PPAR-γ, and GAPDH. (D) Relative expression of inflammation-associated microRNAs miR-21 and miR-30d-5p. Data are presented as mean ± SD (n = 5 per group). iWAT: interscapular white adipose tissue; SED: sedentary; HIIT: high-intensity interval training. * p < 0.05 versus SED; ** p ≤ 0.001 versus SED.

At the protein level, quantitative densitometry confirmed these findings, showing a 55% reduction in NLRP3 protein expression ([95% CI -0.75 to -0.35], p < 0.001, η^2^ = 0.835; Fig. 3-Ci) and a 135% increase in PPAR-γ protein expression ([95% CI 0.28 to 2.43], p = 0.02, η^2^ = 0.514; Fig. 3-Cii). Representative Western blot images are shown in Fig. 3-Ciii.

At the post-transcriptional level, HIIT significantly reduced miR-21 expression by 51% ([95% CI -0.80 to - 0.21], p = 0.004, η^2^ = 0.657) and miR-30d-5p expression by 57% ([95% CI -0.88 to -0.28], p = 0.002, η^2^ = 0.716) compared to the SED group, indicating epigenetic regulation of adipose tissue inflammatory signaling (Fig. 3-D).

## 4. Discussion

The present study investigated the effects of eight weeks of HIIT on epigenetic, gene, and protein markers of inflammatory regulation in iWAT of male Wistar rats. Our findings demonstrate that HIIT significantly reduced body weight and iWAT mass, suppressed the expression of pro-inflammatory mediators, enhanced anti-inflammatory signaling, and downregulated inflammation-associated microRNAs. Together, these results suggest that HIIT induces coordinated adaptations at physiological, transcriptional, protein, and epigenetic levels that collectively improve the inflammatory status of adipose tissue.

The reduction in body weight and adipose tissue mass observed in the present study suggests that HIIT effectively counteracts inactivity-induced adiposity and its associated metabolic dysfunction and chronic low-grad inflammation. Sedentary behavior is well known to promote unfavorable changes in body composition, including increased fat mass and reduced lean mass, which are closely linked to systemic and tissue-specific inflammation (2, 4). White adipose tissue serves a major source of pro-inflammatory cytokines such as TNF-α and IL-6. Enlargement of adipocytes and infiltration of immune cells into adipose tissue activate inflammatory signaling cascades, which in turn promote the development of insulin resistance and increase the risk of chronic diseases, including type 2 diabetes, and immune dysfunction (2, 4, 30).

Regular exercise is recognized as an effective strategy to counteract the adverse effect of sedentary lifestyle (16, 17, 30). Consistent with our findings, previous studies have shown that various exercise modalities, including endurance, resistance, and voluntary exercise, are effective in reducing adiposity and improving inflammatory profiles in rodent adipose tissue (31, 32). Reductions in adipose tissue mass induced by exercise training are closely linked to improved metabolic function. Smaller adipocytes secrete fewer deleterious adipokines, exhibit enhanced insulin sensitivity in target tissues, and contribute to improved energy balance (33, 34). Twelve weeks of swimming exercise has been shown to significantly reduce fat mass and adipocyte size in rats and is accompanied by key cellular and molecular adaptations in adipose tissue, including enhanced insulin receptor density and improved glucose metabolism (35). In summary, the reduction in adiposity observed following exercise training may represent a key mechanism underlying its beneficial effects on adipose tissue inflammation and overall immunometabolic health.

At the molecular level, the present study demonstrated that HIIT significantly reduced NLRP3 expression at both gene and protein levels in iWAT. NLRP3 is a key intracellular receptor that forms the inflammasome complex, leading to caspase-1 activation and subsequent maturation of IL-1β and IL-18, which amplify inflammatory responses and impair insulin signaling (7). The observed reduction in NLRP3 expression is consistent with previous reports indicating that regular aerobic training attenuates NLRP3 expression and associated inflammatory mediators in adipose tissue, vasculature, and immune cells, thereby lowering systemic inflammation and oxidative stress (36). For example, eight weeks moderate-intensity aerobic training performed at 12-20 m/min for 20-50 min/day (∼70% VO_2_ max) has been shown to reduce NLRP3 protein expression in adipose tissue of high-fat diet (HFD)-fed C57BL/6J mice (37). Taken together, these findings suggest that downregulation of NLRP3 may contribute to attenuation of inflammasome activity and improvement of adipose tissue inflammatory status following HIIT.

In addition, our results showed that HIIT significantly decreased TNF-α mRNA levels, another key inflammatory mediator. TNF-α, a well-known pro-inflammatory cytokine, plays a central role in adipose tissue inflammation and metabolic dysfunction by impairing insulin signaling, activating NF-κB and JNK pathways, and reducing adiponectin expression (38). Previous studies have reported inconsistent effects of exercise training on TNF-α expression. In contrast to our findings, twelve weeks of MICT, performed at 50% VO_2_ max with a frequency of 5 days per week, appeared insufficient to alter adipokine expression, including adiponectin, IL-6, and TNF-α in subcutaneous abdominal adipose tissue, or their circulating levels in obese women (39), whereas twelve weeks of voluntary running (15 ± 3 km/ week) significantly increased TNF-α expression at both gene and protein levels in mesenteric fat of insulin-resistant rats fed a high-sucrose diet (40). Conversely, HIIT has been shown to markedly reduce TNF-α expression in visceral adipose tissue of HFD-fed rats following a protocol consisting of five repeated 2-min treadmill sprints performed at 80–90% VO_2_ max interspersed with 1-min active recovery at 30–35% VO_2_max (41). Similar findings have been reported in aged rats, where HIIT consisting of repeated 4-min running bouts at 15 m/min (45–55% VO_2_ max) followed by 1-min sprints at 25 m/min (90–95% VO_2_ max) for nine repetitions reduced TNF-α expression in perirenal adipose tissue (42). Consistent with these findings, the present study showed that HIIT led to a significant reduction in TNF-α gene expression in iWAT of rats. These discrepancies may be attributed not only to differences in exercise intensity and modality, but also to variations in intervention duration, total exercise volume, species differences (humans versus rodents), and tissue-specific responses to exercise.

Alongside these reductions in inflammatory mediators, the present study demonstrated a significant upregulation of PPAR-γ expression in iWAT following HIIT. PPAR-γ is a nuclear receptor and transcription factor that regulates lipid metabolism, insulin sensitivity, and adipocyte differentiation, while also exerting strong anti-inflammatory effects via inhibition of NF-κB signaling (6). Our findings are consistent with previous evidence showing that eight weeks of HIIT, consisting of repeated 4-min low-speed running at 15 m/min (45–55% VO_2_ max), followed by 1-min high-speed running at 25 m/min (90–95% VO_2_ max) at 0% incline for nine repetitions, upregulated both PPAR-γ gene and protein expression in perirenal adipose tissue of rats (42). Similarly, MICT (55-65% of VO_2_ max) has been reported to increase PPAR-γ gene expression in mesenteric fat, accompanied by improvement in adipocyte function and insulin sensitivity (43). Taken together, these findings suggest that exercise training promotes a shift toward an anti-inflammatory and metabolically favorable adipose tissue phenotype. This effect may be mediated through PPAR-γ-dependent regulation of lipid metabolism and inflammatory pathways, thereby contributing to improved adipose tissue function and insulin sensitivity.

We also found that HIIT significantly increased IL-10 gene expression in iWAT. IL-10 is an anti-inflammatory cytokine that suppresses excessive immune activation by inhibiting pro-inflammatory cytokine production and supporting insulin sensitivity (43, 44). The increase in IL-10 observed in the present study is in line with previous findings showing that eight weeks of MICT, performed on a treadmill at ∼ 55–65% VO_2_ max, enhances IL-10 expression in adipose tissue of rodents (43, 45). In addition, a recent study demonstrated that ten weeks of HIIT, consisting of five repeated 2-minute treadmill sprints performed at 80–90% VO_2_ max interspersed with 1-minute active recovery at 30–35% VO_2_ max, upregulated IL-10 expression in visceral adipose tissue and reduced insulin resistance in HFD-fed Wistar rats (41). Collectively, these findings indicate that different exercise modalities, including HIIT, enhance IL-10-mediated anti-inflammatory signaling, thereby contributing to improved adipose tissue immunometabolic function.

In addition to transcriptional and protein-level changes, the present study also examined microRNAs as epigenetic regulators of adipose tissue function. MicroRNAs are small non-coding RNAs that regulate gene expression at the post-transcriptional level by inhibiting mRNA translation or promoting mRNA degradation (46).

Notably, miR-21 is a key regulator of inflammatory and metabolic processes, was downregulated by HIIT in the present study. Usually, the elevated expression of miR-21 is associated with activation of inflammatory pathways and suppression of anti-inflammatory responses (11, 12, 47) and observed in conditions such as obesity, type 2 diabetes, and immune disorders . In adipose tissue of obese and diabetic individuals, miR-21 has been implicated in insulin resistance and chronic inflammation (12). Recent evidence suggests that regular physical activity modulates miR-21 in metabolically active tissues, leading to improvements in insulin sensitivity and mitochondrial function (48, 49). In animal models of type 2 diabetes, an eight-week aerobic exercise program performed at 50–60% of VO_2_ max, combined with magnesium supplementation markedly suppressed miR-21 expression in visceral adipose tissue. These molecular adaptations were accompanied by reductions in fasting glucose and insulin, improvement of the adipokine profile, and attenuation of inflammatory signaling, indicating a broader impact on systemic metabolic homeostasis (48). Our results extend these findings by demonstrating that HIIT induced a significant reduction in miR-21 expression in iWAT, highlighting HIIT as an alternative exercise modality in the regulation of miR-21.

Moreover, the present study demonstrated that HIIT downregulated miR-30d-5p. miRNA-30d is implicated in diverse biological processes, including adipocyte differentiation, glucose metabolism, and immune regulations (13-15). Its isoform, miR-30d-5p, derived from the 5′ arm of pre-miR-30d (50), plays a critical role in regulating genes associated with inflammation, cellular differentiation, and tumorigenesis (15). Elevated miR-30d expression has been observed in visceral adipose tissue of patients with type 2 diabetes, suggesting a potential role in metabolic dysregulation (13). Recent evidence indicates that miR-30d-5p modulates inflammatory responses in metabolically active tissues and cardiac tissue following exercise (15, 49). For example, eight weeks of resistance exercise attenuated the pathological upregulation of miR-30d-5p in type 2 diabetic mice. This downregulation was associated with enhanced insulin signaling and improvements in glucose and lipid metabolism. Mechanistically, the beneficial effects were mediated through activation of the SIRT1/PGC1-α axis, enhancing mitochondrial biogenesis and dynamics in skeletal muscle (49). Collectively, these findings suggest that HIIT-induced downregulation of miR-30d-5p may play a key role in improving adipose tissue immunometabolic function.

Despite these promising findings, several limitations should be considered. The present study focused on a selected set of molecular markers involved in inflammatory regulation within adipose tissue; however, a broader range of additional inflammatory and immunometabolic mediators may also contribute to these responses and warrant future investigation. Future studies should therefore explore additional mechanisms, including immune cell infiltration, upstream signaling pathways, and mitochondrial adaptations. In addition, although the HIIT protocol has previously been shown to reduce fasting plasma glucose and enhance insulin sensitivity in rats, suggesting that the observed molecular adaptations likely occur concomitantly with improvements in metabolic health, fasting plasma glucose and insulin sensitivity were not directly assessed in the present study.Furthermore, investigating whether these findings translate to human populations will be essential for determining the clinical relevance of HIIT-induced molecular adaptations. Identifying the downstream targets of miR-21 and miR-30d-5p in adipose tissue may also provide deeper insight into the epigenetic mechanisms through which exercise regulates inflammation and metabolic function.

In conclusion, eight weeks of HIIT induced significant physiological, molecular, and epigenetic adaptations in adipose tissue of rats. The combined reduction in adiposity, suppression of inflammatory mediators, and modulation of key microRNAs suggests that HIIT improves adipose tissue function through coordinated regulation of multiple biological pathways. These findings further support the role of HIIT as an effective and time-efficient intervention for mitigating obesity-related inflammation and promoting metabolic health.

## Abbreviations

HFD: High Fat Diet
HIIT: High-Intensity Interval Training
IL: Interleukin
iWAT: Interscapular White Adipose Tissue
MICT: Moderate-Intensity Continuous Training
miR: microRNA
NLRP3: NOD-like receptor protein 3
PPAR-γ: Peroxisome Proliferator-Activated Receptor Gamma
RT-qPCR: Reverse Transcription Quantitative PCR
SED: Sedentary
TNF-α: Tumor Necrosis Factor-Alpha
WAT: White Adipose Tissue
WB: Western Blot

## Data

The datasets generated and analyzed during the current study are available from the corresponding authors upon reasonable request.

## Generative AI

The authors used Google Gemini solely for creation of the schematic illustration depicting the exercise protocol (Figure 1). The authors reviewed, verified and edited the figure as necessary and take full responsibility for its accuracy and content. No generative AI tools were used in the analysis, interpretation of data, or preparation of the scientific conclusions presented in this manuscript.

